# Measuring and modeling diffuse scattering in protein X-ray crystallography

**DOI:** 10.1101/033746

**Authors:** Andrew H. Van Benschoten, Lin Liu, Ana Gonzalez, Aaron S. Brewster, Nicholas K. Sauter, James S. Fraser, Michael E. Wall

## Abstract

X-ray diffraction has the potential to provide rich information about the structural dynamics of macromolecules. To realize this potential, both Bragg scattering, which is currently used to derive macromolecular structures, and diffuse scattering, which reports on correlations in charge density variations must be measured. Until now measurement of diffuse scattering from protein crystals has been scarce, due to the extra effort of collecting diffuse data. Here, we present three-dimensional measurements of diffuse intensity collected from crystals of the enzymes cyclophilin A and trypsin. The measurements were obtained from the same X-ray diffraction images as the Bragg data, using best practices for standard data collection. To model the underlying dynamics in a practical way that could be used during structure refinement, we tested Translation-Libration-Screw (TLS), Liquid-Like Motions (LLM), and coarse-grained Normal Modes (NM) models of protein motions. The LLM model provides a global picture of motions and were refined against the diffuse data, while the TLS and NM models provide more detailed and distinct descriptions of atom displacements, and only used information from the Bragg data. Whereas different TLS groupings yielded similar Bragg intensities, they yielded different diffuse intensities, none of which agreed well with the data. In contrast, both the LLM and NM models agreed substantially with the diffuse data. These results demonstrate a realistic path to increase the number of diffuse datasets available to the wider biosciences community and indicate that NM-based refinement can generate dynamics-inspired structural models that simultaneously agree with both Bragg and diffuse scattering.

**Significance:** The structural details of protein motions are critical to understanding many biological processes, but they are often hidden to conventional biophysical techniques. Diffuse X-ray scattering can reveal details of the correlated movements between atoms; however, the data collection historically has required extra effort and dedicated experimental protocols. We have measured three-dimensional diffuse intensities in X-ray diffraction from CypA and trypsin crystals using standard crystallographic data collection techniques. Analysis of the resulting data is consistent with the protein motions resembling diffusion in a liquid or vibrations of a soft solid. Our results show that using diffuse scattering to model protein motions can become a component of routine crystallographic analysis through the extension of commonplace methods.

## Introduction

X-ray crystallography can be a key tool for elucidating the structural basis of protein motions that play critical roles in enzymatic reactions, protein-protein interactions and signaling cascades (van den Bedem and Fraser, 2015). X-ray diffraction yields an ensemble-averaged picture of the protein structure: each photon simultaneously probes multiple unit cells that can vary due to internal rearrangements or changes to the crystal lattice. Bragg analysis of X-ray diffraction only yields the mean charge density of the unit cell, however, which fundamentally limits the information that can be obtained about protein dynamics (Clarage and Phillips, 1997; Keen and Goodwin, 2015).

A key limitation inherent in Bragg analysis is that alternative models with different correlations between atomic motions can yield the same mean charge density (Kuzmanic et al., 2011). The traditional approach to modeling atom movement is to assume a single structural model with individual atomic displacement parameters (B factors). Given sufficient data, anisotropic displacement factors can be modeled, yielding directional insights into motions that might cause variations in the crystal. When the data are more limited, Translation-Libration-Screw (TLS) structural refinement, in which motions are described using rigid body segments of the molecule (Schomaker and Trueblood, 1968), has emerged as a common tool to model protein domain movements in crystallography and has been used by 22% of PDB depositions (Painter and Merritt, 2005, 2006). However, TLS refinements that vary in the rigid body definitions can predict very different motions while maintaining equivalent agreement to Bragg X-ray diffraction data (Urzhumtsev et al., 2015; Van Benschoten et al., 2015).

Additional sources of information have been used to overcome the inherent limitations of Bragg analysis in identifying collective protein motion. Patterns of steric clashes between alternative local conformations (van den Bedem et al., 2013) or time-averaged ensemble refinement (Burnley et al., 2012) can be used to suggest certain modes of concerted motion. However, the atomistic details of these correlated motions may only be reliably (yet indirectly) identified at high resolution, and time-averaged ensemble refinement is additionally complicated by the use of an underlying TLS model to account for crystal packing variations (Burnley et al., 2012). Alternative methods such as solid-state NMR experiments (Ma et al., 2015) or long time scale molecular dynamics simulations (Janowski et al., 2013; Janowski et al., 2015; Wall et al., 2014b) can be used to probe the structural basis of crystal packing variations and internal protein motions.

Complementary information about internal protein motions also can be obtained in the X-ray crystallography experiment itself by analysis of diffuse scattering. Diffuse scattering arises when deviations away from a perfect crystal cause X-rays to be diffracted away from Bragg reflections. When the deviations are due to crystal vibrations, they can be described using textbook temperature diffuse scattering theory (see, e.g. (James, 1948)). When each unit cell varies independently, the diffuse intensity is proportional to the variance in the unit cell structure factor (Guinier, 1963) which is equivalent to the Fourier transform of the Patterson function of the charge density variations. The approximation of independent unit cells can break down when correlations extend across unit cell boundaries; however, motions with long correlation lengths result in diffuse intensity concentrated in the immediate neighborhood of Bragg peaks. When analyzing the more broadly distributed diffuse intensity that corresponds to small correlation lengths (Caspar et al., 1988; Clarage et al., 1992; Wall et al., 1997a; Wall et al., 1997b) the contribution of inter-unit cell atom pairs is a small fraction of the total signal, which is therefore dominated by internal protein motions.

Several approaches have been used to connect macromolecular diffuse scattering data to models of protein motion and lattice disorder. Notably, Peter Moore has emphasized the need to validate TLS models using diffuse scattering (Moore, 2009), as has been performed in a limited number of cases (Doucet and Benoit, 1987; Perez et al., 1996; Van Benschoten et al., 2015). Good agreement with the data has previously been observed for liquid-like motions (LLM) models (Caspar et al., 1988; Clarage et al., 1992; Wall et al., 1997a; Wall et al., 1997b). In the LLM model, the atoms in the protein are assumed to move randomly, like in a homogeneous medium; the motions were termed "liquid-like” by Caspar et al (Caspar et al., 1988) because the correlations in the displacements were assumed to fall off exponentially with the distance between atoms. Like the LLM model, normal modes (NM) models treat the protein as a softer substance than the TLS model while still treating it as a solid. Unlike the LLM model, however, normal-mode analysis (NMA) provides a much more detailed picture of the conformational ensemble, enabling a more direct connection to putative mechanisms of protein function (Yang et al., 2007). In addition, like TLS refinement, the normal modes refinement methods that have been developed for Bragg analysis use few additional parameters (Gniewek et al., 2012; Kidera et al., 1994; Lu and Ma, 2008; Ni et al., 2009; Poon et al., 2007). Unfortunately these programs are not currently available in the standard builds of the major refinement software. Reasonable qualitative agreement previously been seen using normal modes to model diffuse intensity in individual diffraction images (Faure et al., 1994; Mizuguchi et al., 1994), and, more recently, the fit of alternative coarse-grained elastic network models to diffuse scattering data of staphylococcal nuclease has been investigated (Riccardi et al., 2010).

There is also a longstanding interest both in using diffuse scattering to validate improvements in MD simulations and in using MD to derive a structural basis for the protein motions that give rise to diffuse scattering (Clarage et al., 1995; Faure et al., 1994; Héry et al., 1998; Meinhold et al., 2007; Meinhold and Smith, 2005a, 2005b, 2007; Wall et al., 2014b). Recent advances in computing now enable microsecond duration simulations (Wall et al., 2014b) that can overcome past barriers to accurate calculations seen using 10 ns or shorter MD trajectories (Clarage et al., 1995; Meinhold and Smith, 2005a).

Despite the fact that diffuse scattering analysis is relatively well developed in small-molecule crystallography (Welberry, 2004) and materials science (Keen and Goodwin, 2015), it has been underutilized in protein crystallography. There are relatively few examples of diffuse data analyzed using individual diffraction images from protein crystallography experiments, including studies of tropomyosin (Chacko and Phillips, 1992; Phillips et al., 1980), 6-phosphogluconate dehydrogenase (Helliwell et al., 1986), yeast initiator tRNA (Kolatkar et al., 1994), insulin (Caspar et al., 1988), lysozyme (Clarage et al., 1992; Doucet and Benoit, 1987; Faure et al., 1994; Mizuguchi et al., 1994; Perez et al., 1996), myoglobin (Clarage et al., 1995), Gag protein (Welberry et al., 2011), and the 70s ribosome subunit (Polikanov and Moore, 2015). Moreover, there are an even smaller number of examples involving complete three-dimensional diffuse data sets; these include studies of staphylococcal nuclease (Wall et al., 1997b), and calmodulin (Wall et al., 1997a).

To exploit the increased information that is potentially available from diffuse scattering, there is a pressing need to increase the number of proteins for which complete three-dimensional diffuse datasets have been experimentally measured. Conventional data collection procedures use oscillation exposures to estimate the full Bragg intensities. In contrast, the complete three-dimensional datasets measured by Wall *et al*. (Wall et al., 1997a; Wall et al., 1997b) used specialized methods for integrating three-dimensional diffuse data from still diffraction images. Similar methods now can be generalized and applied to other systems using modern beamlines and X-ray detectors. In particular, the recent commercial development of pixel-array detectors (PADs), which possess tight point-spread functions and single-photon sensitivity (Gruner, 2012), have created new opportunities for measuring diffuse scattering as a routine tool in protein crystallography experiments using more conventional data collection protocols.

Here, we present diffuse scattering datasets for the human proline isomerase cyclophilin A (CypA) and the bovine serine protease trypsin. These datasets substantially increase the amount of experimental three-dimensional diffuse scattering data available to the macromolecular crystallography community, providing a necessary foundation for further advancement of the field (Wall et al., 2014a). To assess the potential for routine collection of diffuse datasets in crystallography, rather than expending a great deal of effort in optimizing the diffuse data and collecting still images (Wall et al., 1997a; Wall et al., 1997b), we used oscillation images obtained using best practices for high-quality Bragg data collection. The resulting datasets are of sufficient quality that the diffuse scattering can discriminate among alternative TLS refinements (Van Benschoten et al., 2015), Liquid-Like Motion (LLM) models (Caspar et al., 1988; Clarage et al., 1992), and Normal Modes (NM) models ((Meinhold and Smith, 2007; Mizuguchi et al., 1994; Riccardi et al., 2010)). Moreover, the agreement of the NM models with both Bragg and diffuse scattering data suggests a path forward for anisotropic refinement of atomic displacements, while maintaining a small set of parameters, using both data sources simultaneously. Our results demonstrate that diffuse intensity can, and should, be measured in a typical X-ray crystallography experiment and indicate that diffuse X-ray scattering can be applied broadly as a tool to understand the conformational dynamics of macromolecules.

## Results

### Experimental diffuse data show crystallographic symmetry

The symmetrized anisotropic diffuse datasets processed by LUNUS (**Figure 1**) are shown in **Figure 2A** (CypA) and **Figure 2D** (trypsin) and are available as supplementary material. The CypA dataset is 98% complete to a resolution of 1.4 Å, while the trypsin map is 95% complete to 1.25 Å resolution. We used the Friedel symmetry and Laue group symmetry to quantify the level of crystallographic symmetry in each anisotropic map. To evaluate the degree Friedel symmetry, we averaged intensities between Friedel pairs to create a symmetrized map *I_Friedel_* and calculated the Pearson Correlation Coefficient (PCC) between the symmetrized and unsymmetrized data to obtain the statistic CC_Friedel_. For CypA and trypsin, CC_Friedel_ = 0.90 and 0.95 respectively, demonstrating that diffuse intensities obey Friedel symmetry. To assess the degree of Laue group symmetry, we averaged P222-related reflections (the Laue symmetry corresponding to the P 21 21 21 space group of both CypA and trypsin crystals) to produce the symmetrized intensities, *I*_P222_. The linear correlation CC_Sym_ was then computed between the symmetrized and unsymmetrized intensities. The correlations were substantial for both CypA (CC_Sym_ = 0.70) and trypsin (CC_Sym_ = 0.69). Thus, our data are consistent with the diffuse intensity following the Bragg peak symmetry. The trypsin data were integrated using one degree oscillation frames, while the CypA data were integrated using 0.5 degree oscillation frames. The comparable degree of symmetry in the CypA and trypsin data suggests that the measurement of diffuse intensity is robust with respect to this difference in data collection.

**Figure 1.**
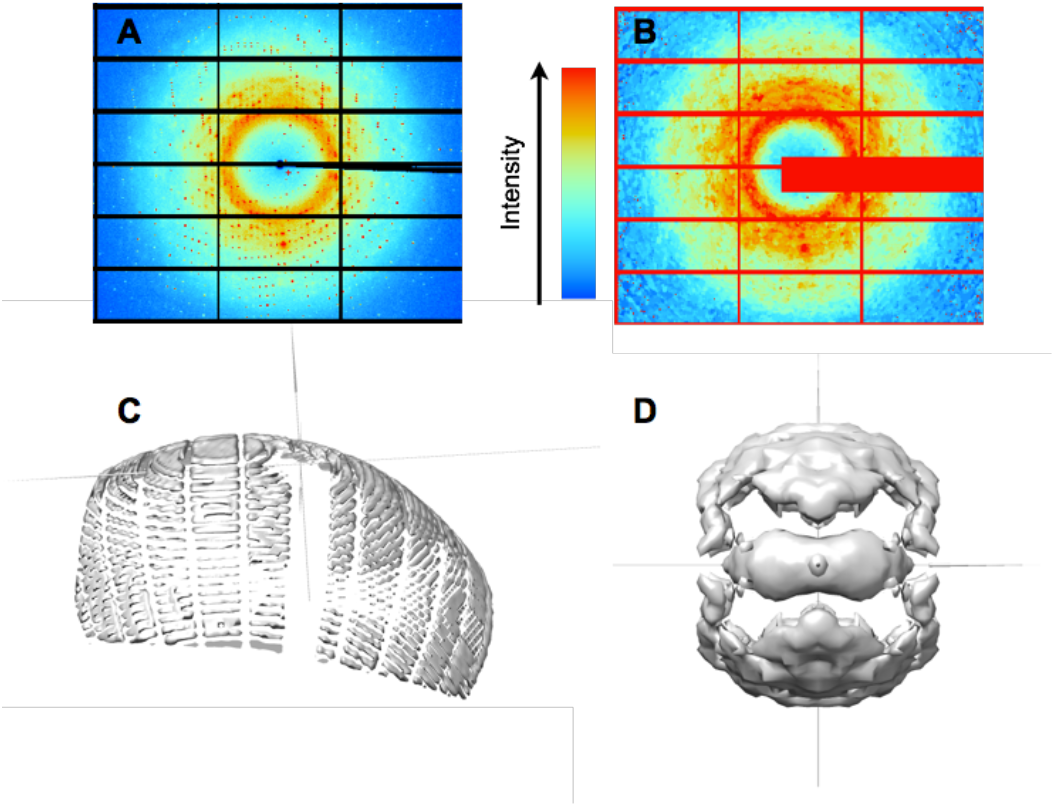
Steps in diffuse data integration. (A) Raw CypA diffraction images are processed (B) to remove Bragg peaks and enable direct comparisons of pixel values to models. (C) Pixels in diffraction images are mapped to reciprocal space and values of diffuse intensity are accumulated on a three-dimensional lattice; each diffraction image produces measurements of diffuse intensity on the surface of an Ewald sphere. (D) The data from individual images is combined and symmetrized to yield a nearly complete dataset (isosurface at a value of 65 photon counts in the total intensity, before subtracting the isotropic component).

**Figure 2.**
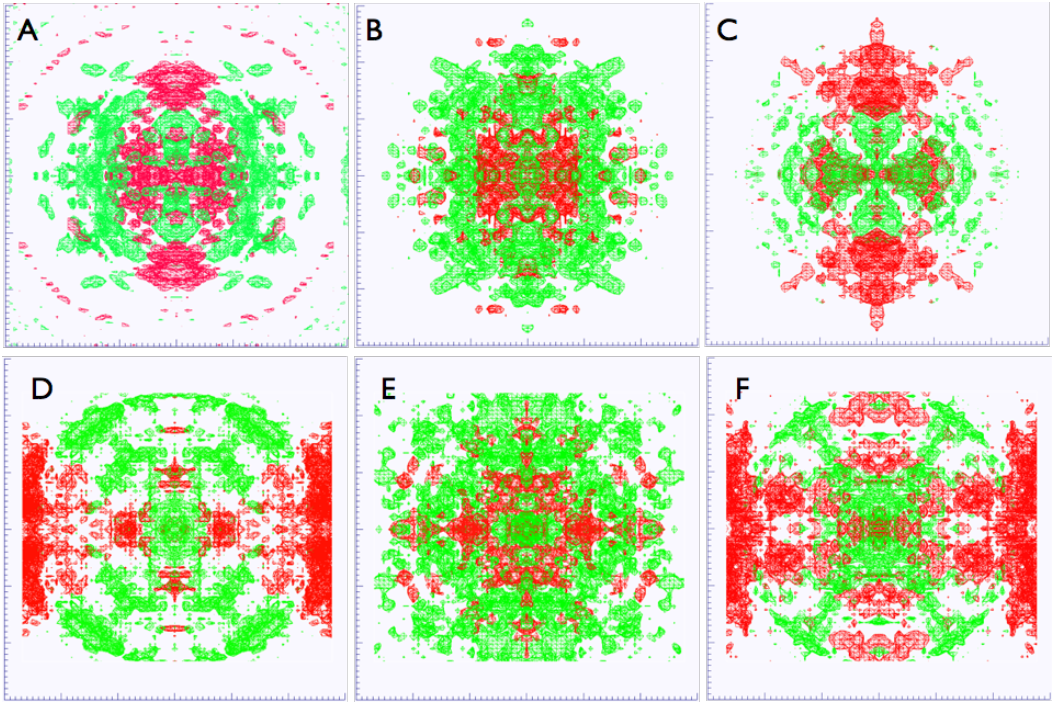
Visualization of anisotropic diffuse intensities. (A) CypA experimental data with isosurfaces shown using wireframes at a level of 2 photon counts in the resolution range 4.16 Å – 2.97 Å. Positive intensity is rendered in green, negative in red. (B) Isosurfaces for diffuse scattering predicted by the CypA LLM model. (C) Residual diffuse scattering (experimental data (A) minus LLM (B)). (D) Trypsin experimental data with isosurfaces shown using wireframes at a level of 3 photon counts in the resolution range 4.53 Å – 3.26. (E) Isosurfaces for diffuse scattering predicted by the Trypsin LLM model. (F) Residual diffuse scattering (experimental data (D) minus LLM (E)).

### TLS models yield low correlation with diffuse scattering data

To investigate how well TLS models agree with the molecular motions in the CypA crystal, we compared the experimental diffuse data to intensities calculated from three alternative TLS models: *phenix, tlsmd* and *whole molecule* (**Figure 3A-D**). Although all three models predict different motions, the R-factors are very similar: R,R-free = 16.4%,18.1% for the *whole molecule* and *Phenix* models; and 16.2%,18.1% for the TLSMD model. The correlations between the calculated diffuse intensity for these models and the anisotropic experimental data are low: 0.03 for the *phenix* model; 0.04 for the *TLSMD* model; and 0.14 for the *whole molecule* model. In addition, the pairwise correlations of the calculated diffuse intensities are low: 0.066 for *whole molecule/TLSMD;* 0.116 for *whole molecule/Phenix;* and 0.220 for *Phenix/TLSMD*.

**Figure 3.**
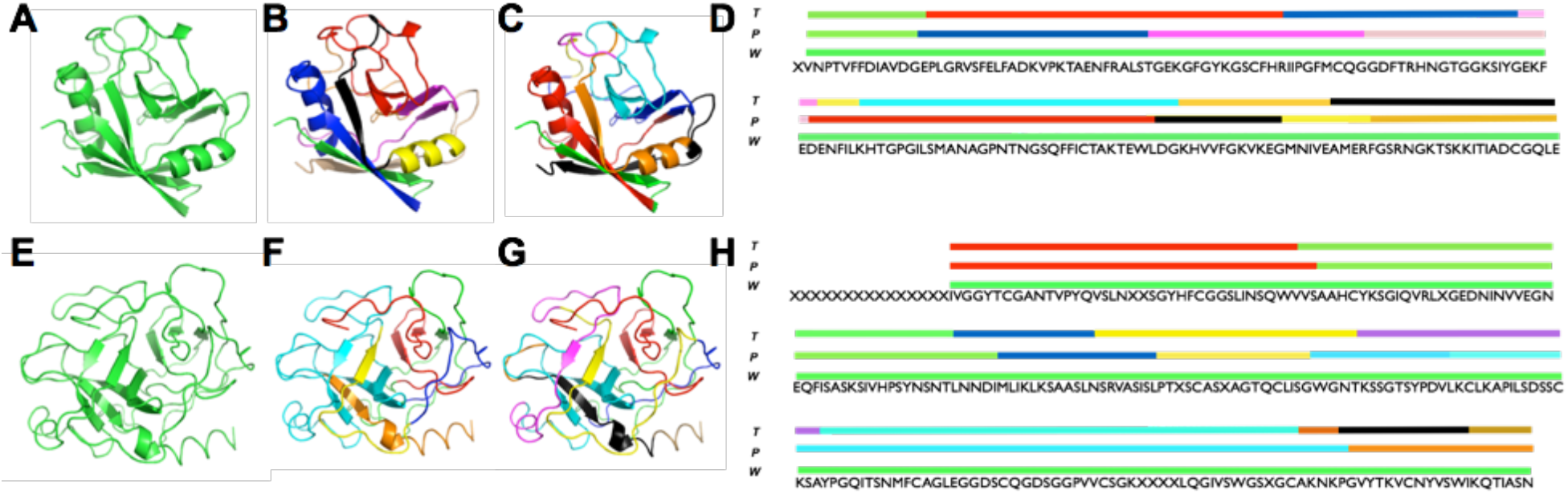
Rigid body domain definitions used for TLS models. CypA and Trypsin TLS groups shown on the tertiary structure for whole molecule (A, E), Phenix (B, F), and TLSMD (C, G) and shown on the primary sequence (D, H).

Like CypA, the three trypsin TLS models (**Figure 3E-H**) yielded very similar R,R-free values: 15.1%,16.7% for the *whole molecule* model; 15.3%,16.6% for the Phenix model; and 15.2%,16.6% for the TLSMD model. Correlations between the calculated and experimental diffuse intensities are again low: 0.02 for the *Phenix* and *TLSMD* models, and 0.08 for the *Whole molecule* model. Comparisons of the calculated anisotropic diffuse intensity show that the *Whole molecule* motion is dissimilar to both the *Phenix* and *TLSMD* predictions (PCC = 0.03 and 0.05, respectively). In contrast, the *Phenix* and *TLSMD* models yield much more similar diffuse intensities (PCC = 0.515). The relatively high correlation between these models is consistent with the similarity in the TLS groups (**Figure 3F-H**).

There are several possible explanations for the low correlation between the TLS model and diffuse data for CypA and trypsin. First, TLS domain groupings other than those identified here might yield higher agreement with the data. Second, the method used for generating ensembles (Van Benschoten, 2015) assumes that TLS domains vary independently; it is possible that accounting for correlations among the domains would more accurately describe the variations. Lastly, similar to the rigid body motions model of Doucet & Benoit (Doucet and Benoit, 1987), the correlations among TLS domains might lead to substantial correlations across unit cell boundaries, which would produce small scale diffuse features in the immediate neighborhood of Bragg peaks. The data integration methods used here cannot resolve these features, as the measurements are mapped to a Bragg lattice. Methods to integrate the small-scale features in protein crystallography onto a finer three-dimensional reciprocal space grid do exist (Wall et al., 1997a) and might be used to address this last possibility in the future. In any case, the low correlation of TLS models with the diffuse intensity for CypA and trypsin suggests that the variations in the protein crystal might not be best explained by motions of relatively large, rigid domains, and instead might involve motions that are correlated on a shorter length scale than accounted for by these models.

### Liquid-like motions models yield substantial correlation with diffuse scattering data

One model that accounts for short-range correlations is Liquid-Like Motions (LLM) (Caspar et al., 1988; Clarage et al., 1992). The LLM model assumes that atomic displacements are uncorrelated between different unit cells, but are correlated within the unit cells. The correlation in the displacements is assumed to decay exponentially as *f(x) = e^−x/Y^*, where x is the separation of the atoms, and **γ** is the length scale of the correlation. The displacements of all atoms are assigned a standard deviation of **σ**. The LLM model previously has been refined against three-dimensional diffuse intensities obtained from crystalline staphylococcal nuclease (Wall et al., 1997b) and calmodulin (Wall et al., 1997a), yielding insights into correlated motions.

We refined isotropic LLM models of motions in CypA and trypsin against the experimental diffuse intensities (**Figure 2B, E and Methods**). The CypA model was refined using data in the resolution range 31.2 Å – 1.45 Å, and the trypsin model using 68 Å – 1.46 Å data. For CypA, the refinement yielded **γ** = 7.1 Å and **σ** = 0.38 Å with a correlation of 0.518 between the calculated and experimental anisotropic intensities. The highest correlation between data and experiment occurs in the range 3.67 Å – 3.28 Å, where the value is 0.74 (**Figure 4A**). For the trypsin dataset, the refinement yielded **γ** = 8.35 Å and **σ** = 0.32Å with a correlation of 0.44, which is lower than for CypA. The peak value is 0.72 in the resolution range 4.53 Å -4.00 Å (**Figure 4B**).

**Figure 4.**
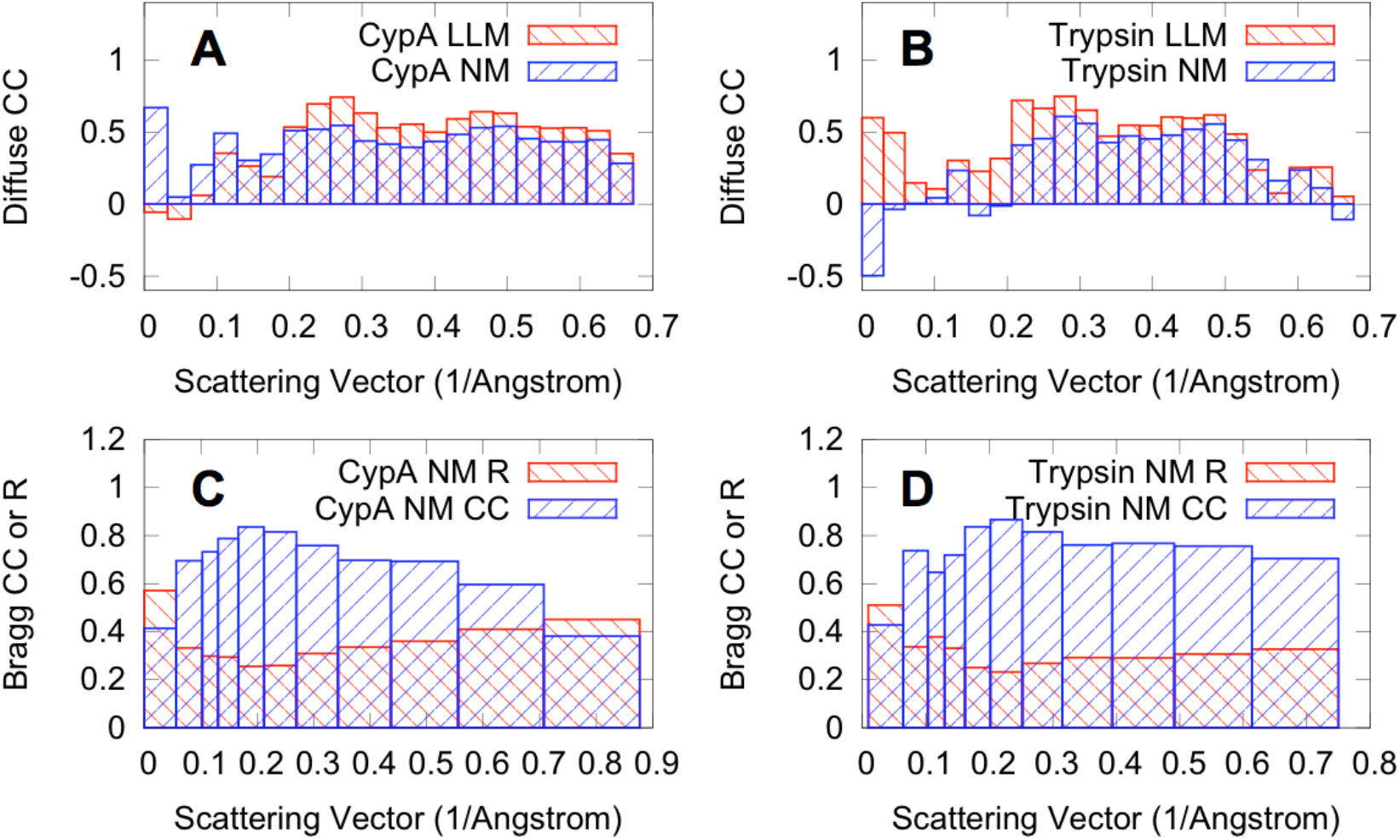
Agreement of models of protein motions with diffuse and Bragg data. (A, B) Linear correlation coefficients (CC) between diffuse data and LLM (red bars) or NM models (blue bars) computed by resolution shell for (A) CypA and (B) Trypsin. (C, D) Correlations and R-factors between Bragg data and NM models computed by resolution shell for (C) CypA and (D) Trypsin. Agreement factors for the diffuse and Bragg data were computed using LUNUS (Wall, 2009) and *Phenix* (Adams et al., 2010), respectively.

The refined LLM models also were compared to the data using simulated diffraction images. Images corresponding to frame number 67 of the CypA data were obtained using the LLM model (Fig. 5A) and the integrated diffuse data (Fig. 5B). The main bright features above and below the origin are similar between the two. Many of the weaker features also appear to be similar, both at high and low resolution. The similarity is diminished but still apparent for images obtained for frame number 45 of trypsin (Supplementary Fig. S1). These simulations provide a visual confirmation of the substantial correlations obtained for the three-dimensional diffuse intensity.

**Figure 5.**
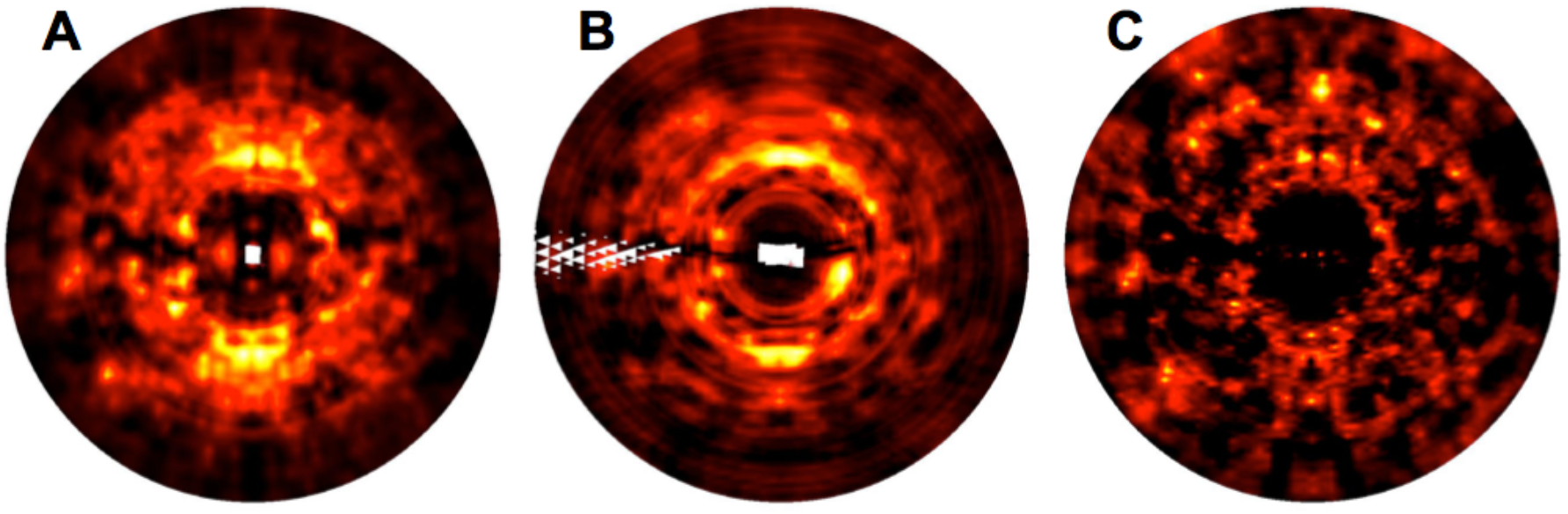
Simulated diffraction images for CypA frame 67 obtained using: (A) liquid-like motions model; (B) integrated 3D diffuse data; (C) elastic network model. Lighter colors correspond to stronger intensity. White regions correspond to pixel values where there are missing values in the corresponding three-dimensional lattice (Methods).

The substantial correlation of the LLM model with the diffuse data for CypA and trypsin indicates that the variations in the protein crystal can be approximately described using a model of the protein as a soft, homogeneous medium. The model implies that the motions of atoms separated by more than 7-8 Å are relatively independent, and that atoms that are closer to each other move in a more concerted way.

### Normal-modes can model both diffuse and Bragg scattering data

To assess the potential of NMA to be developed for diffuse scattering studies, we developed coarse-grained elastic network models of the CypA and trypsin unit cells (Methods). The C_α_ coordinates and B-factors for the NM models are by definition identical to those derived from the Bragg data (Methods). To assess the agreement of specific NM-derived conformational variations with the Bragg data, we generated 50 member ensembles from the Hessian (Methods). We applied a single isotropic B-factor across all atoms ensuring that all deviations in the individual magnitude and anisotropy originated from the normal modes (Fig. S2). The correlations were high across resolution shells (Fig. 4 C, D), yielding overall R-factors of 38% (CypA) and 31% (Trypsin) (Tables S1 and S2). We also calculated the predicted diffuse intensity from the NM models: the correlation of the CypA model with the data is 0.41 in the resolution range 31.2 Å – 1.45 Å, and the correlation of the trypsin model with the data is 0.38 in the resolution range 68 Å – 1.46 Å. The agreement with the data is substantial within individual resolution shells (Fig. 4). The NM simulated diffraction image for CypA (Fig. 5C) shows bright features that are found in the data (Figs. 5B). The relative strength at high versus low-resolution is higher than in the data, however, suggesting that this NM model is too rigid; this discrepancy might be addressed by softening the intra-residue interactions and optimizing the model against the diffuse scattering data directly. The comparisons of simulated diffraction images for trypsin are consistent with the findings for CypA (Fig. S1).

Overall the agreement of the NM models with the data assessed using either 3D diffuse scattering datasets (Fig. 4) or simulated diffraction images (Figs. 5, S1) is substantial but slightly less than for the LLM models. However, it is important to interpret this comparison in light of the fact that the covariance matrices of the NM models were normalized to agree with the Bragg data and not parameterized against the diffuse data (Methods), while the LLM model is parameterized against the diffuse data. The agreement with the Bragg data is currently limited by the fact that the parameter optimization used only the refined C_α_ positions and B-factors to agree with the Bragg data and that heteroatoms, such as solvent, were not included in the calculations. Collectively, these results point to the potential for normal modes to be refined jointly against Bragg and diffuse scattering data as an alternative atomic displacement model, replacing TLS or individual B-factors.

## Discussion

Diffuse X-ray scattering is a potentially valuable yet little exploited source of information about macromolecular dynamics. Diffuse intensities can double the total number of measured data points in the crystallographic experiment while providing a parallel dataset against which structural dynamical models can be refined or validated. Until now measurement of three-dimensional diffuse scattering data only has been pursued in dedicated efforts requiring extra still diffraction images and substantial optimization of experimental design. The present collection of two new datasets obtained using oscillation images using best current practices in room-temperature protein crystallography (Fraser et al., 2011), and the use of the data in evaluating TLS, LLM, and NM models, illustrates the potential for using diffuse scattering to increase understanding of protein structure variations in any X-ray crystallography experiment, representing a significant step towards moving diffuse scattering analysis into the mainstream of structural biology.

Diffuse data obtained for CypA and trypsin can distinguish among the TLS, LLM, and NM models of motions. However, the agreement with the data is somewhat lower than in previous LLM models of three-dimensional diffuse scattering (Wall et al., 1997a; Wall et al., 1997b). In this study, the correlation of the LLM model with the data was 0.518 in the range 31.2 Å – 1.45 Å for CypA, and 0.44 in the range 68 Å – 1.46 Å for trypsin; in comparison, the correlation was 0.595 in the range 10 Å – 2.5 Å for staphylococcal nuclease (Wall et al., 1997b) and 0.55 in the range 7.5 Å – 2.1 Å for calmodulin (Wall et al., 1997a). Some possible explanations for the lower agreement for CypA and trypsin include: the use of higher resolution data in the present studies; that LLM might be a better description of motions in staphylococcal nuclease and calmodulin than in CypA and trypsin; and that the measurements might have been more accurate in the past experiments, as the data collection was tailored for diffuse scattering. The apparent alignment of the residual intensity distribution with the unit cell axes (**Figures 2C, 2F**) also suggests that an anisotropic LLM model might be more appropriate than an isotropic LLM model for CypA and trypsin.

The agreement of the LLM models with three-dimensional experimental diffuse data across multiple systems warrants further consideration for using diffuse scattering in model refinement and validation. A key finding is that the agreement of the LLM models with the diffuse data is higher than the TLS models, which currently are used widely in protein crystallography.

Interestingly, the 7-8 Å length scale of the correlations is comparable to the size of the TLS domains; however, compared to the sharp domains of the TLS model, the exponential form of the correlations indicates that there is a smooth spatial transition between the correlated and uncorrelated atoms in the LLM. The smooth transition might be key to the increased agreement of the LLM with the diffuse data compared to the rigidly defined regions of the TLS model.

Our findings also support the use of NM models in combination with diffuse scattering for model refinement and validation. The exploratory work here, which did not use diffuse data for parameterization, indicates that NM models contain features that can capture aspects of the diffuse scattering data, and motivates further work to incorporate NM in both Bragg and diffuse refinement. In the coarse-grained NM model in Eq. (2), the residues are treated as rigid, which artificially increases anisotropic features at high resolution. We performed a limited exploration of models with decreased intra-residue atom correlations: so far these models have led to lower correlations with the data; however, in principle such models should be more accurate. Future work will focus on developing computationally efficient methods for optimizing the accuracy of the coarse-grained NM models and for using all-atom NM to model diffuse scattering.

Overall, the three-dimensional diffuse scattering data obtained here for CypA and trypsin, and previously for staphylococcal nuclease (Wall et al., 1997b) and calmodulin (Wall et al., 1997a) suggest that the protein structure varies more like a soft material than like a collection of independent rigid domains. An important consideration in developing these new refinement methods is to maintain a key advantage of TLS refinement at lower resolutions: the introduction of relatively few new parameters for refinement. This requirement also would be satisfied by NMA, which has a low computational cost and general applicability, making it a promising model for integrating diffuse scattering into crystallographic model building and refinement (Wall et al., 2014a). Whether this pursuit is well-motivated hinges on whether new biological insights can be gained from atomic displacements generated by NM models refined against Bragg and diffuse data. Indeed, although use of TLS in model refinement is now widespread, it scarcely has been used to generate biological hypotheses (for exceptions, see: (Chaudhry et al., 2004; Henzler-Wildman et al., 2007)). In contrast to TLS models, elastic network NM models have been widely used to draw functional inferences (Bahar et al., 2010). Both the encouraging agreement of the NM models with the diffuse scattering and the potential for NM models to yield new insights about the importance of conformational dynamics in protein function provide a strong motivation for further developing NM models for protein X-ray crystallography.

Diffuse scattering also can be used to validate models of molecular motions other than those considered here, including models produced by ensemble refinement (Burnley et al., 2012); multiconformer modeling performed by discrete (Keedy et al., 2015; van den Bedem et al., 2009) or continuous (Burling and Brünger, 1994; Kuriyan et al., 1991; Wall et al., 1997a) conformational sampling; and molecular dynamics simulations (Clarage and Phillips, 1994; Clarage et al., 1995; Faure et al., 1994; Héry et al., 1998; Janowski et al., 2013; Janowski et al., 2015; Meinhold et al., 2007; Meinhold and Smith, 2005a, 2005b; Wall et al., 2014b). In particular, molecular dynamics simulations now provide sufficient sampling to yield robust calculations of diffuse intensity (Wall et al., 2014b), and these can be used to consider a myriad of intramolecular motions (e.g., loop openings and side chain flips) (Wilson, 2013) and lattice dynamics. Polikanov and Moore (Polikanov and Moore, 2015) recently have demonstrated the importance of lattice vibrations in explaining experimental diffuse scattering measurements of ribosome crystals, which indicates that models should simultaneously account for correlations that are coupled both within and across unit cell boundaries (Clarage et al., 1992; Wall et al., 1997a); accounting for lattice vibrations more accurately also might yield improved Bragg integration (Wall et al., 2014a). Moreover, comparisons of crystal simulations and diffuse scattering can provide a new observable for benchmarking improvements in energy functions and sampling schemes (Janowski et al., 2015).

Although the initial successes of dynamics-based models of diffuse scattering indicates that crystal defects can play a secondary role in contributing to the diffuse signal, at least in some cases, consideration of crystal defects might become important to achieve the highest model accuracy and most general applicability of diffuse scattering in crystallography. Additionally, as more X-ray data from both brighter conventional and XFEL light sources, accounting for all sources of Bragg and diffuse scattering will be necessary to model the total scattering needed for innovative phasing applications (Gaffney and Chapman, 2007). In summary, the new datasets presented here demonstrate that diffuse scattering can now be routinely collected and that using these data will help us obtain an increasingly realistic picture of motion in protein crystals, including integrated descriptions of intramolecular motions, lattice vibrations, and crystal defects.

## Methods

### Protein purification and crystallization

Trypsin crystals were obtained according to the method of Liebschner *et.al* (Liebschner et al., 2013). Lyophilized bovine pancreas trypsin was purchased from Sigma-Aldrich (T1005) and dissolved at a concentration of 30 mg/mL into 30mM HEPES pH 7.0, 5 mg/mL benzamidine and 3mM CaCl_2_. Crystals were obtained from a solution of 200mM Ammonium sulfate, 100mM Na cacodylate pH 6.5, 20% PEG 8000 and 15% glycerol. CypA was purified and crystallized as previously described (Fraser et al., 2009). Briefly, the protein was concentrated to 60 mg/mL in 20mM HEPES pH 7.5, 100mM NaCl and 500mM TCEP. Trays were set with a precipitant solution of 100mM HEPES pH 7.5, 22% PEG 3350 and 5mM TCEP. Both crystal forms were obtained using the hanging-drop method.

### Crystallographic data collection

Diffraction data were collected on beamline 11-1 at the Stanford Synchrotron Radiation Lightsource (Menlo Park, CA). X-ray diffraction images were obtained using a Dectris PILATUS 6M Pixel Array Detector (PAD). Each dataset was collected from a single crystal at an ambient temperature of 273K. To prevent dehydration, crystals were coated in a thin film of paratone with minimal surrounding mother liquor. For CypA, a single set of 0.5 degree oscillation images were collected and used for both Bragg and diffuse data processing. A total of 360 images were collected across a 180 degree phi rotation. The Trypsin diffraction data consisted of one degree oscillations across a 135 degree phi rotation; this dataset was similarly used for both Bragg and diffuse data analysis. Both datasets were collected to optimize the Bragg signal, not the diffuse signal. Although not used here, we note that data collection using a PAD with fine phi slicing should be especially well suited for simultaneous collection of Bragg and diffuse data, as it would enable integration of diffuse intensity at a tunable level of detail in reciprocal space.

### Bragg data processing

Bragg diffraction data were processed using XDS and XSCALE (Kabsch, 2010) within the *xia2* software package (Winter et al., 2013). Molecular replacement solutions were found using Phaser (McCoy et al., 2007) within the *Phenix* software suite (Adams et al., 2010). The PDB search models were 4I8G for trypsin, and 2CPL for CypA. Initial structural refinement was performed using *phenix.refine* (Afonine et al., 2012). The strategy included refinement of individual atomic coordinates and water picking. Both the X-ray/atomic displacement parameters and X-ray/stereochemistry weights were optimized. Isotropic B-factors were chosen for the initial structures to allow for non-negligible R-factor optimization by subsequent TLS refinement strategies. All structures were refined for a total of 5 macrocycles. Statistics for these initial crystal structure models are shown in **Table 1**.

**Table 1.** Refinement statistics for CypA and trypsin models, before TLS modeling is applied.

|  | CypA | Trypsin |
| --- | --- | --- |
| Resolution range, Å | 38.66-1.4 | 23.29-1.25 |
| Space group | P 21 21 21 | P 21 21 21 |
| Unit cell, Å | 42.91, 52.44, 89.12 | 54.81, 58.51, 67.42 |
| Completeness (%) | 98 | 95 |
| R<sub>work</sub> (%) | 17.88 | 15.9 |
| R<sub>free</sub> (%) | 19.5 | 17.41 |
| RMS (bonds, Å) | 0.007 | 0.013 |
| RMS (angles, degrees) | 1.16 | 1.61 |
| Ramachandran favored % | 97 | 98 |
| Ramachandran allowed % | 3 | 2 |
| Ramachandran outliers % | 0 | 0 |
| Clashscore | 0.79 | 2.59 |
| Average B-factor, Å<sup>2</sup> | 21.42 | 14.57 |

### Diffuse data integration

An overview of the diffuse data integration process is presented in **Figure 1**. Image processing was performed using the LUNUS collection of diffuse scattering tools (Wall, 2009). Pixels corresponding to the beam stop and image edges were masked using the *punchim* and *windim* methods. To focus on the diffuse intensity, which compared to Bragg peaks has low individual pixel values (while, being more broadly distributed in reciprocal space, having comparable total integrated intensity), pixel values outside of the range 1-10,000 photon counts were masked using *threshim*. The beam polarization was determined by analyzing the first frame to determine the azimuthal intensity profile within a 100 pixel wide annulus about the origin, and by fitting the resulting profile to the theoretical profile (Wall, 1996). Pixel values then were corrected for beam polarization using *polarim*. A solid-angle normalization (*normim*) correction was also applied. Mode filtering was used to remove Bragg peaks from diffraction images. This was accomplished using *modeim*, with a mask width of 20 pixels and a single bin for each photon count increment. These steps produced diffraction images in which pixel values could be directly compared to model diffuse intensities. This procedure, originally developed for experiments on staphylococcal nuclease (Wall et al., 1997b), is similar to the steps used by Polikanov and Moore (Polikanov and Moore, 2015) to process individual ribosome diffraction images for analysis of diffuse scattering data.

The Lunus processed frames were used to integrate the diffuse data onto a 3D lattice. The integration was performed using a python script that calls DIALS methods within the Computational Crystallography Toolbox (CCTBX; (Grosse-Kunstleve et al., 2002; Parkhurst et al., 2014)). The script obtains an indexing solution using the *real_space_grid_search* method and uses the results to map each pixel in each diffraction image to fractional Miller indices *h’k’l’* in reciprocal space. It sums the intensities from pixels in the neighborhood of each integer Miller index *hkl* and tracks the corresponding pixel counts, while ignoring pixels that fall within a ½ X ½ X ½ region about *hkl*. It writes the intensity sums and pixel counts for each frame on a grid, populated on an Ewald sphere that varies according to the crystal orientation for each image (**Figure 1C**). A radial scattering vector intensity profile was calculated for each frame using the Lunus *avgrim* method and was used to scale diffuse frames across the entire dataset. The Lunus *sumlt* and *divlt* methods were used to compute the mean diffuse intensity at each grid point using the scaled sums and pixel counts from all of the frames.

Because the model diffuse intensities were computed without considering solvent, experimental and model diffuse intensities were compared using just the anisotropic component of the signal, which is primarily due to the protein (Wall et al., 2014b). The Lunus *avgrlt* and *subrflt* routines were applied to subtract the radial average and obtain the anisotropic signal. Signal intensities were then symmetrized using *phenix.reflection_file_converter* to obtain a dataset for comparison to models. Datasets were compared to each other and to models using linear correlations computed using the *phenix.reflection_statistics* tool.

All images are available on SBGrid Data Grid (https://data.sbgrid.org/dataset/68/ for CypA; https://data.sbgrid.org/dataset/201/ for Trypsin) and the integrated diffuse scattering maps are available as Supplementary Material.

### Simulated diffraction images

Diffuse scattering images were simulated using methods similar to those for data integration in the previous section. After finding an indexing solution, a frame corresponding to the desired simulated image was selected from the data set. This frame was used as a template for obtaining a mapping of each pixel to fractional Miller indices. The new value of each pixel was obtained by linear interpolation of the values of diffuse intensity between the nearest-neighbor integer points *hkl* for which diffuse intensity was either measured (as in the previous section) or calculated (as in below sections on liquid-like motions and normal modes models). In the case of the synthetic images computed from the diffuse data, the images greatly enhanced the ability to visualize diffuse features compared to the original diffraction images (Supplementary Figs. S3, S4); the enhancement is due to the improved statistics obtained by averaging many pixel values to obtain a measurement at each value of *hkl*. The simulated images were processed to enhance visualization of diffuse features: the minimum pixel value was computed within each pixel-width annulus about the beam center, and was subtracted from each pixel value within the annulus. Images were displayed using Adxv (Arvai, 2012), with display parameters selected for meaningful comparison of the diffuse features.

### TLS structure refinement and diffuse scattering model

Three independent TLS refinements were performed for CypA (**Figure 3A-D**). The *Whole molecule* selection consists of the entire molecule as a single TLS group. The *Phenix* selection consists of the 8 groups (residues 2-14, 15-41, 42-64, 65-84, 85-122, 123-135, 136-145 and 146-165) identified by *phenix.find_tls_groups*. The *TLSMD* selection consists of 8 groups (residues 2-15, 16-55, 56-80, 81-85, 86-91, 92-124, 125-143 and 144-165) identified by the TLS Motion Determination web server (Painter and Merritt, 2005, 2006). All TLS refinement was performed within *phenix.refine* through 5 macrocycles. Aside from the inclusion of TLS refinement, these macrocycles were identical to the initial structure refinement described above.

Similarly, for trypsin, we selected Whole *Molecule, Phenix*, and *TLSMD* TLS refinement strategies as described above (**Figure 3E-H**). The *Phenix* selection consists of 7 TLS groups: residues 16-54, 55-103, 104-123, 124-140, 141-155, 156-225 and 226-245. The *TLSMD* selection consists of 9 groups: residues 16-52, 53-98, 99-115, 116-144, 145-171, 172-220, 221-224, 225-237 and 238-245.

Structural ensembles of the CypA and trypsin TLS motions were generated through the *Phenix.tls_as_xyz* method (Urzhumtsev et al., 2015). Each ensemble consisted of 1,000 random samples of the underlying TLS atomic displacement distributions, assuming independent distributions for each domain. Diffuse scattering models were calculated from the TLS ensembles using *Phenix.diffuse* (Van Benschoten et al., 2015). CypA and trypsin models were generated to a final resolution of 1.2 Å and 1.4 Å respectively, to match the resolution of the experimental data.

### Liquid-like motions model

We computed Liquid-like motions (LLM) models of diffuse scattering using the structures refined prior to the TLS refinements (CypA: PDB; Trypsin: PDB). For both CypA and trypsin, the temperature factors for all atoms were set to zero and squared calculated structure factors *I_0_(hkl)* were computed using the *structure_factors, as_intensity_array*, and *expand_to_p1* methods in CCTBX (Grosse-Kunstleve et al., 2002; Parkhurst et al., 2014). The Lunus *symlt* method was used to fill in missing values in reciprocal space using the appropriate P222 Laue symmetry.

Given a correlation length γ and amplitude of motion σ, the diffuse intensity predicted by the LLM model was calculated as

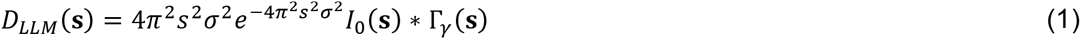

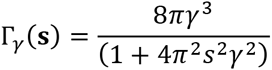

Fourier methods in Lunus (*fftlt*) were used to compute the convolution. The agreement with the data was quantified by computing a linear correlation as a target function, using the anisotropic intensities (**Diffuse data integration**). Optimization of the target with respect to γ and σ was performed in a python script using *scipy.optimize* (www.scipy.org) with the Powell minimization method.

### Normal modes model

The diffuse intensity was computed using a normal modes (NM) model of correlated atom displacements, using methods similar to Riccardi et al. (Riccardi et al., 2010). Atomic coordinates and isotropic displacement parameters were obtained from PDB entries 5F66 (CypA) and 5F6M (trypsin) and were parsed and expanded to the P1 unit cell using the iotbx.pdb methods in CCTBX (Grosse-Kunstleve et al., 2002). The Hessian matrix **H** was defined using a modified anisotropic elastic network model (Atilgan et al., 2001), with springs between C_α_ atoms (*i,j*) within a cutoff radius of 25 Å. The spring force constants were computed as *ke^−r_ij_/λ^*, where *r_ij_* is the closest distance between atoms *i* and *j*, either in the same unit cell or in neighboring unit cells; *λ* = 10.5 Å; and *k* = 1 for *r_ij_* < 25 Å and *k* = 0 otherwise (the nonzero value of *k* is arbitrary due to the normalization employed below). Covariances of atom pair displacements *v_ij_* = 〈**r**_i_ ˙ **r**_j_〉 were obtained using the pseudoinverse of **H** as described in (Atilgan et al., 2001). The values of *v_ij_* were renormalized to *ϕ_ij_* = *v_ij_ σ_i_σ_j_/(v_ii_ v_jj_)*^1/2^ using the isotropic displacement parameters σ_i_ of the *i*^th^ C_α_ atom from the Bragg refinement; the model was thus consistent with the refined crystal structure.

The diffuse intensity was computed as

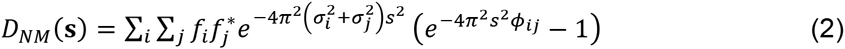

where *f_i_* is the structure factor of the combined atoms in the residue associated with the *i*^th^ C_α_ atom. Structure factors were computed using a two-gaussian approximation of atomic form factors; the parameters were obtained using the eltbx.xray_scattering methods in CCTBX (Grosse-Kunstleve et al., 2002); phase factors were applied using the atomic coordinates.

The Bragg intensities were computed from ensembles generated by using the first 10 nonzero eigenvectors of **H** with corresponding inverse eigenvalues as their weights. Because the overall scale of the spring constant was arbitrary in the NM model (see above), the amplitudes of motion were too large using the absolute eigenvalues; they were therefore scaled to maintain the connectivity of the backbones. 50 member ensemble models were generated by Normal Mode Wizard (*NMWiz*) (Bakan et al., 2011), which is a *VMD* (Humphrey et al., 1996) plugin. A single B-factor of 10 was applied to all atoms in the ensemble. Structure factors were generated using *phenix.fmodel* and compared to the experimental data using *phenix.reflection_statistics* (Adams et al., 2010).

## Acknowledgments

We thank Pavel Afonine for computational assistance in converting and comparing structure factors. We are grateful to the UC Office of the President, Multicampus Research Programs and Initiatives grant MR-15-338599 and the Program for Breakthrough Biomedical Research, which is partially funded by the Sandler Foundation. Use of the Stanford Synchrotron Radiation Lightsource, SLAC National Accelerator Laboratory, is supported by the U.S. Department of Energy, Office of Science, Office of Basic Energy Sciences under Contract No. DE-AC02- 76SF00515. The SSRL Structural Molecular Biology Program is supported by the DOE Office of Biological and Environmental Research, and by the National Institutes of Health, National Institute of General Medical Sciences (including P41GM103393). N.K.S. was supported by NIH grant GM095887. J.S.F. was supported by a Searle Scholar Award from the Kinship Foundation, a Pew Scholar Award from the Pew Charitable Trusts, a Packard Fellowship from the David and Lucile Packard Foundation, NIH OD009180, NIH GM110580, and NSF STC-1231306. M.E.W. was supported by the US Department of Energy under Contract DE-AC52-06NA25396 through the Laboratory-Directed Research and Development Program at Los Alamos National Laboratory (LANL). The LANL technical release number is LA-UR-15-28934.

